# ATF3 drives senescence by reconstructing accessible chromatin profiles

**DOI:** 10.1101/2020.08.13.249458

**Authors:** Chao Zhang, Xuebin Zhang, Yiting Guan, Xiaoke Huang, Lijun Zhang, Xiao-Li Tian, Wei Tao

## Abstract

Chromatin architecture and gene expression profile undergo tremendous reestablishment during senescence. However, the regulatory mechanism between chromatin reconstruction and gene expression in senescence remain elusive. The chromatin accessibility is an excellent perspective to reveal the latent regulatory elements. Thus, we depicted the landscapes of chromatin accessibility and gene expression during HUVECs senescence. We found that chromatin accessibilities are re-distributed during senescence. The senescence related increased accessible regions (IARs) and the decreased accessible regions (DARs) are mainly distributed in distal intergenic regions. The DARs are correlated with the function declines caused by senescence, whereas the IARs are involved in the regulation for senescence program. Moreover, the heterochromatin contributes most of IARs in senescent cells. We identified that the AP-1 transcription factors, especially ATF3 is responsible for driving chromatin accessibility reconstruction in IARs. In particular, DNA methylation is negatively correlated with chromatin accessibility during senescence. AP-1 motifs with low DNA methylation may improve their binding affinity in IARs and further opens the chromatin nearby. Our results described a dynamic landscape of chromatin accessibility whose remodeling contributes to the senescence program. And we identified a cellular senescence regulator, AP-1, which promotes senescence through organizing the accessibility profile in IARs.

## Introduction

Cellular senescence is known as an irreversible cell-cycle arrest further bringing a series of progressive cellular states and phenotypic changes(van Deursen, 2014). Senescent cells play intricate roles in many biological processes including tumor suppression, embryogenesis, tissue repair, host immunity, aging and age-related disorders(Chou & Effros, 2013; He & Sharpless, 2017; Jun & Lau, 2010; Munoz-Espin et al., 2013; Reddel, 2009; Storer et al., 2013). The mechanisms of senescence have historically been viewed as replicated senescence and stress induced senescence(de Magalhaes & Passos, 2018; Hayflick & Moorhead, 1961; Robert H. te Poele, 2002; Serrano, Lin, McCurrach, Beach, & Lowe, 1997). Although the effects of senescence have been known well, the jury is still out on the upstream of regulatory network. In many candidate hallmarks that may contribute to aging(Lopez-Otin, Blasco, Partridge, Serrano, & Kroemer, 2013), epigenetic alterations including spontaneous or passive changes are pretty compelling.

The epigenome serves as a bridge connecting the genotype with the phenotype which is responsible to interpret the information of genome(Pal & Tyler, 2016). Epigenetics has been considered an extremely important contributor to senescence and aging(Fraga et al., 2005; Payel Sen, Shah, Nativio, & Berger, 2016). DNA methylation patterns, posttranslational modification of histones and chromatin remodeling are well regarded primarily epigenetic alterations. Demethylation of whole genome was taken place while local hypermethylation still exist during replicative senescence(Hazel A. Cruickshanks et al., 2013; Wilson & Jones, 1983). DNA methylation patterns can be used to estimate the state of cellular senescence which is referred as DNA methylation clock(Field et al., 2018; Franzen et al., 2017; Horvath, 2013). Histone modifications reflect the different status of chromatin structures and we can investigate the regulatory mechanism of senescence and aging based on the varying patterns of histone modifications. For examples, H3K9me3 is reduced in Werner syndrome model cells (W. Zhang et al., 2015), decrease of H4K16ac is observed in Zmpste-24-null fibroblasts(Krishnan et al., 2011), loss of H4K20me3 promotes senescence and aging (Lyu et al., 2018).

The alterations of DNA methylation and histones modifications bring changes to chromatin architecture which impresses the profiles of gene expression and further determines the cell fate. It has been concerned several decades ago that chromatin structure changes have close connection with senescence(Howard, 1996; Macieira-Coelho, 1980, 1991). There exists a chromatin remodeling procedure in cell nucleus during senescence which involves the formation of senescence-associated heterochromatin foci (SAHF) in where specific proliferation-associated genes are repressed(Narita et al., 2006; Narita et al., 2003; R. Zhang et al., 2005). While the local facultative heterochromatin is constructed, there is a global decrease of constitutive heterochromatin in senescence that is hypothesized as “heterochromatin loss model of aging”(Tsurumi & Li, 2012; Villeponteau, 1997). After years of exploration, many senescence associated chromatin remodeling factors come into the mind of the people. Both silence and overexpression of brahma-related gene 1 (Brg1) which is the ATPase subunit of SWI/SNF complex could induce the cellular senescence in human or rat mesenchymal stem cells (MSCs) (Alessio et al., 2010; Napolitano et al., 2007). Loss of the components of NURD complex leads to ageing-related chromatin defects(Pegoraro et al., 2009). Recently, a RNAi screening of epigenetic proteins indicated that p300 drives senescence mediated by *de novo* super enhancer formation(P. Sen et al., 2019). These researches show the close correlation between rearrangement of chromatin architecture and senescence. We can illustrate the mechanism of senescence and aging from the top view of chromatin landscape.

As a hallmark of active DNA regulatory elements such as enhancer and promoter, chromatin accessibility plays the role in regulating gene expression. The redistribution of accessible chromatin reflects the dynamic physical interactions between chromatin-binding factors and DNA, which cooperatively regulate gene expression profile(Klemm, Shipony, & Greenleaf, 2019). The analyses of chromatin accessibility reveal the dynamic gene regulatory networks in various physiological and pathological processes(Cao et al., 2018; Corces et al., 2018; Gao et al., 2018; John et al., 2011; L. Liu et al., 2019; Y. Liu et al., 2018; Xie et al., 2020). These researches indicate that chromatin accessibility may be a potential factor to predetermine the stimuli response program and even cell fate. AP-1 transcription factors including ATF, Fos and Jun participate in a wide range of cellular program like proliferation and apoptosis (Ameyar, Wisniewska, & Weitzman, 2003; Karin, Liu, & Zandi, 1997; Shaulian & Karin, 2001). More and more evidences indicated that AP-1 is responsible for establishment and maintenance of open chromatin orchestrating with gene expression in multiple cell types which partially function as a pioneer factor (Biddie et al., 2011; Britton et al., 2017; Kurachi et al., 2014; Vierbuchen et al., 2017).

Here, we described the landscape of chromatin accessibility during HUVECs senescence using ATAC-seq which is a powerful tool to reveal the genome-wide chromatin accessibility based on hyperactive Tn5 transposase(J. D. Buenrostro, Giresi, Zaba, Chang, & Greenleaf, 2013; Jason D. Buenrostro et al., 2015). Meanwhile, we integrated the open chromatin landscape and gene expression profile to study the gene regulatory network during senescence. We found that the gene expression and chromatin accessibility undergo reprogramming in senescence. There are senescence-related chromatin regions in which accessibility is increased or decreased gradually during senescence, we named them IARs or DARs respectively. Their functions are quite different in senescence, the IARs related signaling pathways contribute to senescence progression, but the DARs related signaling pathways mainly involve in physiological function. Further analysis showed that the IARs are mostly located in heterochromatin. Moreover, we found DNA methylation levels are negatively correlated with chromatin accessibility in IARs. AP-1 transcription factors were identified to regulate the openness of the IARs. And ATF3, a member of AP-1 family, promoted cellular senescence through reprogramming the chromatin accessibility.

## Results

### Genome-wide mapping of gene expression for replicated senescence

To decode the mechanism of senescence, we construct the replicated senescence system in primary human umbilical vein endothelial cells (HUVECs). The proliferation of HUVECs gradually slows down along with passaging which can be cultured about 40 population doublings (about 20 to 30 passages) *in vitro* (Supplementary Fig. 1A). We defined the 4 passages(P4) as young cells, 14 passages(P14) as middle cells, and 26 passages(P26) as senescent cells according to our experimental conditions. We employed different senescence markers to better evaluate the reliability of our replicated senescence system. The signals of SA-β-Gal and Ki-67 were gradually increased or decreased with the extension of HUVECs culture time, which are classical senescence or proliferation markers(Dimri et al., 1995; Gerdes et al., 1987)(Fig. 1A-C). In addition, the protein level of Lamin B1, a molecular marker of senescence whose protein level decline in most senescence systems was decreased in our system (Fig. 1D). Together, the HUVECs replicated senescence system we established is stable and reliable.

**Fig. 1.**
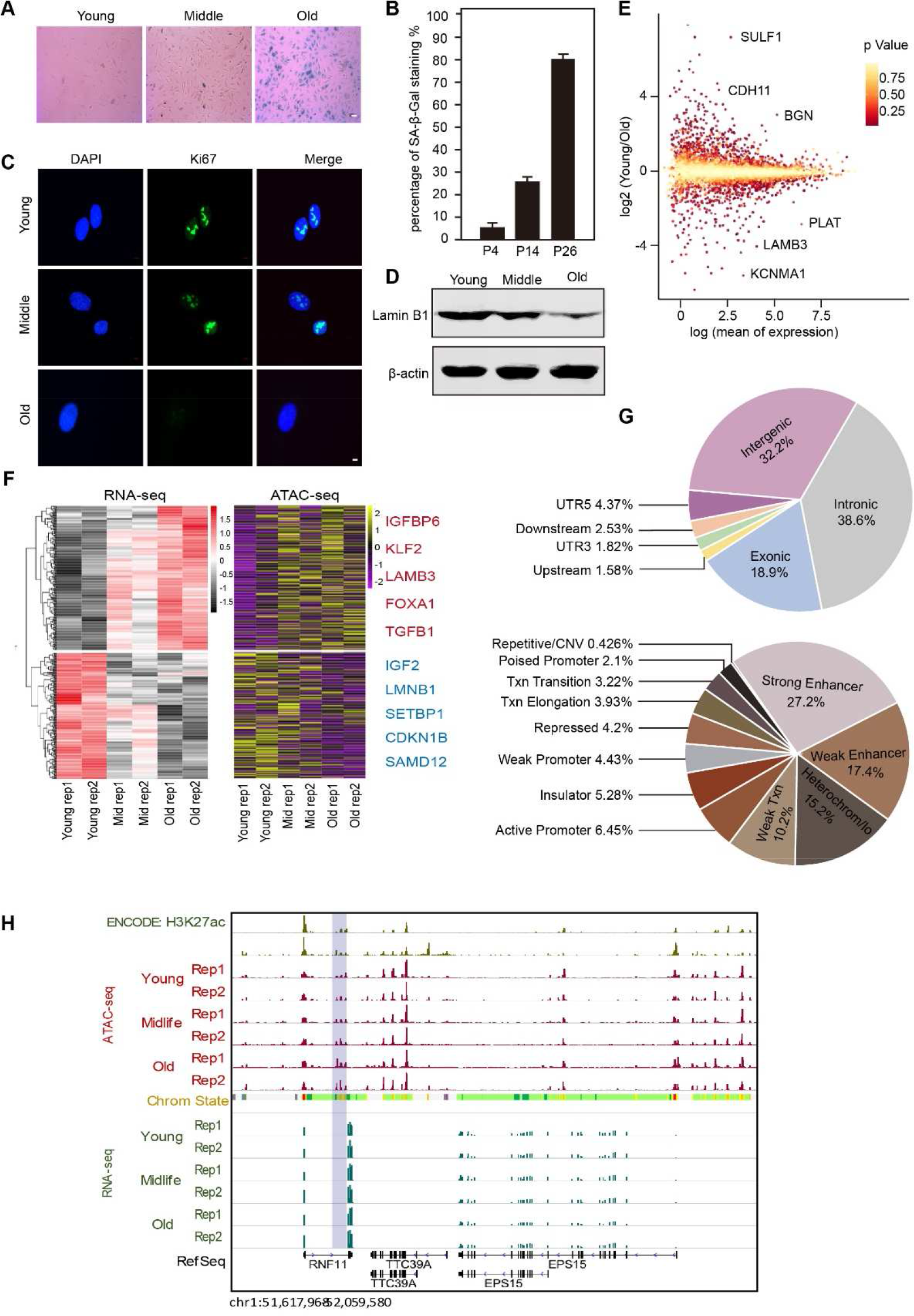
The genome-wide landscape of gene expression and chromatin accessibility for replicated senescence. **A** SA-β-gal staining in young(P4), middel(P14) and old(P26) HUVECs. Scale bar, 50 μm. **B** Diagram of statistics for SA-β-gal positive HUVECs. **C** Immunofluorescence staining of Ki67 in serially passaged HUVECs. Scale bar, 5 μm. **D** Western blot showing the expression of Lamin B1 in HUVECs senescence. β-actin served as loading control. **E** Dispersion plots of RNA-seq signals shows the differentially expressed genes between young and senescent HUVECs. **F** Heatmaps of RNA-seq and ATAC-seq showing the genes which were consistent between expression and accessibility in senescence. The upper part displays the upregulated genes and the lower part displayed the downregulated genes along with senescence. Two independently repeated experiments were performed in young, middle and old HUVECs. **G** Distribution analysis of open chromatins for HUVECs senescence. Upper diagram shows the accessibility analysis for chromatin regions. Lower diagram shows the accessibility analysis for regulatory elements. **H** Snapshot showing the signals of RNA-seq, ATAC-seq and H3K27ac in example regions. Blue shadow shows these signals in the enhancer of RNF11.

Next, to investigate the dynamic changes of gene expression in HUVECs senescence, we performed the RNA-seq and found that the gene expression profile displayed uptrend and downtrend at the same time (Supplementary Fig. 2A and 2B). The expression of CDKN1A(Chang et al., 2000) and TGFBI(B. Li, Wen, Zhao, Tong, & Hei, 2012) gradually increased and the expression of LMNB1(Freund, Laberge, Demaria, & Campisi, 2012) gradually decreased during senescence (Supplementary Fig. 2A-B), which are consistent with previous study. In addition, the expression of some genes rarely reported in aging researches also undergone very significant changes, such as upregulated genes KCNMA1, PLAT, LAMB3, and downregulated genes SULF1, CDH11, BGN, CDKN1B (Fig. 1E and Supplementary Fig. 2), these genes may be related to the physiological functions of HUVECs. In general, the results of RNA-seq also showed that our replicated senescence system is credible. Furthermore, in order to figure out the change patterns of gene expression network in senescence, we performed the GO (Gene Ontology) analysis. The signaling pathways that perturbed greatly during senescence can be roughly classed as three groups, one may be involved in the occurrence and development of senescence, the another may be related with the dysfunction caused by senescence, and the third one may be relevant to functional decline of endothelial cells along with senescence. The up-regulated genes are mainly involved in the PI3K-Akt signaling pathway, which widely responds to external signals to regulate cell growth, but there are some reports that the activation of the PI3K-Akt signaling pathway possibly triggers senescence (Astle et al., 2012) which are consistent with our conclusion. Besides, the focal adhesion signaling pathway and ECM-receptor pathway are also upregulated that are closely related to the formation of senescence phenotypes, such as adhesion plaques and cell migration (Borghesan & O’Loghlen, 2017) (Supplementary Fig. 3A). The DNA replication and cell cycle associated pathways are downregulated which is in line with the definition of cell cycle arrest about senescence (Supplementary Fig. 3B). Moreover, some pathways related to cellular functions were downregulated indicating the function decline caused by senescence. For instances, the thyroid hormone pathway that regulates metabolic homeostasis and the Rap1 pathway that regulates cell adhesion and connection in response to external stimuli (Supplementary Fig. 3B). In summary, our analysis about the gene expression profile and signaling pathways indicated that the senescence system we established is reliable, and revealed the HUVECs senescence pattern that accord with the universal mechanism.

**Fig. 2.**
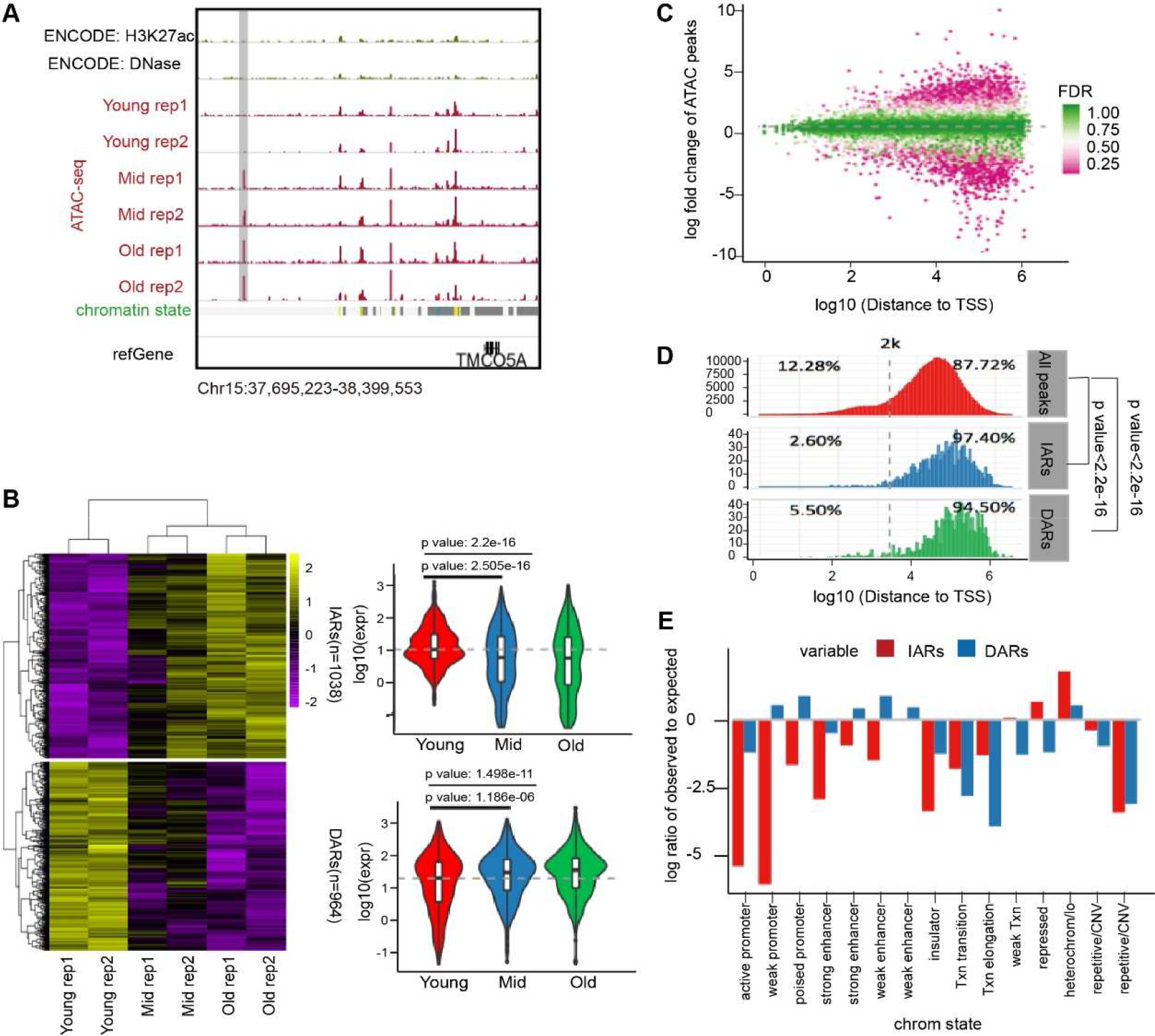
The dynamics of chromatin accessibility during senescence. **A** Snapshot showing the signals distribution of ATAC-seq, DNase-seq and H3K27ac in or near the TMCO5A in HUVECs senescence. Gray shadow shows the gradually increased ATAC-seq signals. **B** The heatmap showing the dynamic remodeling of IARs and DARs during senescence. **C** Scatter diagram showing the distance to TSS of ATAC-seq signals, most of regions significantly changed in senescence are far away from the TSS. **D** Statistics of distribution ratio for IARs and DARs, most of them lie in where are away from TSS. As are significant compared with all peaks. **E** Analysis for the distribution of chromatin states in IARs and DARs.

### The landscape of chromatin accessibility during senescence

Then to explore how the gene expression profile is reconstructed, we performed the ATAC-seq to study the changes of chromatin accessibility during senescence. We first analyzed the reliability of our ATAC-seq data and aligned it with the DNase-seq data of HUVECs from the ENCODE database. The two methods showed similar trends about chromatin accessibility remodeling (Supplementary Fig. 4A). At the same time, we performed the quality checks for our data, the transcription start sites (TSS) possessed the highest enrichment of ATAC-seq signals which is consistent with the reference (Supplementary Fig. 4B)(Klein & Hainer, 2019; Sun, Miao, & Sun, 2019). These results showed that the ATAC-seq data is reliable. We compared the gene expression profile and the chromatin accessibility profile during senescence, and found their change trends are corresponding and unified in senescence. For examples, the significantly upregulated genes IGFBP6(Coppe et al., 2010), KLF2(Taniguchi et al., 2012), LAMB3(Denisenko, Mar, Trawczynski, & Bomsztyk, 2018), FOXA1(Q. Li et al., 2013), TGFB1(Senturk et al., 2010), their chromatin accessibility is also increased remarkably with senescence. The significantly downregulated genes IGF2 (DeChiara, Efstratiadis, & Robertson, 1990), LMNB1, SETBP1(Lucas et al., 2018), CDKN1B(Pruitt, Freeland, Rusiniak, Kunnev, & Cady, 2013), SAMD12, their chromatin accessibility is also decreased remarkably with senescence (Fig. 1F). These results showed that the chromatin accessibility contributed to the expression change of differential genes related to senescence, especially the key genes that trigger senescence.

In order to more clearly show the changes of open chromatin during senescence, we analyzed the distribution characteristics of accessible chromatin regions that significantly remodeled before and after senescence. We found that the intergenic, intronic and exonic regions have the most abundant ATAC-seq signals, and in which the enhancer possessed the most proportion (Fig. 1G and Supplementary Fig. 4C). For example, there are rich ATAC-seq signals in the defined enhancer of RNF11 where the signals of H3K27ac, a marker of chromatin accessibility, are distributed (Fig. 1H). Taken together, these results suggested that we generated high-quality ATAC-seq data in HUVECs replicated senescence, on the base of that, we depicted the landscape of chromatin accessibility and surprisingly found that the ATAC-seq signals of enhancers have the most remarkable changes during senescence.

### The dynamics of chromatin accessibility during senescence

Considering that we have revealed the dynamic landscape of chromatin accessibility during senescence, we wondered investigate whether there are some senescence specific regions that undergone regular changes in chromatin accessibility. We compared the whole-genome ATAC-seq signals at different senescence stages. The number of ATAC-seq signal peaks in young and old cells were more than that in middle (Supplementary Fig. 5A). And we noticed more observed features of ATAC-seq signal peaks in senescence (Supplementary Fig. 5B). Actually, most of signal peaks have no apparent variations while the signal peaks in some specific regions have. Obviously, the statistics of ATAC-seq signal peaks indicated that the chromatin accessibility was rearranged indeed during senescence. And these regions are distributed far away from TSS (Supplementary Fig. 5C). On the basis of the statistics for ATAC-seq signals (Fig. 1G), we also compared the details of chromatin state between different samples from each senescence stage (Supplementary Fig. 5D and 5E). The chromatin regions mainly include enhancers and promoters in which accessibility was remodeled significantly in senescence, and the increased regions are mainly active elements whereas decreased regions are mainly weak elements. Which indicated that the regulation modes among different classes of DNA elements were reconstructed along with senescence. Although there are some differences in the accessibility of different senescence stages in the above analysis, their specific characteristics are not obvious, and they do not show very clear rules. Thus, we need to focus on smaller range to discovery some novel regions.

Next, we focused on some specific chromatin regions that showed a certain remodeling pattern of accessibility during senescence. Surprisingly, we found that there was a type of chromatin regions whose ATAC-seq signal peaks increased consecutively with the progress of senescence, such as the TMCO5, but the H3K27ac and DNase-seq from the database do not have a signal peak here. Thus, some information about senescence can only be found by continuous detection at the time dimension (Fig. 2A). Later, we counted the number of these regions or with similar features to eliminate accidents and confirm that they are indeed characteristic regions in senescence through analyzing whole-genome ATAC-seq peaks. After statistics, we found that there are 1038 regions where the chromatin accessibility gradually increased and 964 regions where the chromatin accessibility gradually decreased during senescence (Fig. 2B). It can be seen that these regions that gradually increased or decreased along with senescence are not special cases. Instead they are specific regions with identified characteristics that may play a unique role in senescence. We defined the former as increased accessible region (IAR) and the latter as decreased accessible region (DAR).

Later, in order to have a more comprehensive understanding of the characteristics of IARs or DARs, and continue to explore whether these regions have practical significance closely related to senescence. We raised three questions for them, the first question is what is their distribution regularity on genome. The second question is which chromatin states corresponding to these regions. The third question is whether these regions can be connected with the senescence program. For the first question, we analyzed the distance between the rearranged accessible chromatin and TSS during senescence. We found that the regions whose accessibility changed the most during senescence are almost far away from TSS (Fig. 2C). After analyzing the distance to TSS of IARs and DARs, we found that the vast majority of IARs (97.4%) and DARs (94.5%) are located far away from the TSS (Fig. 2D). For the second question, we aligned the ATAC-seq signals of IARs and DARs to database to explore the chromatin state information of the IARs and DARs, and found a surprising feature, that is, IARs are mainly distributed in heterochromatin, and DARs are mainly distributed in promoters and enhancers (Fig. 2E). Taken together, our results revealed that the chromatin accessibility profile was rearranged during senescence, and there were some senescence specific regions with regular changes in accessibility, of which IARs are mainly distributed in heterochromatin that already defined and DARs are mainly distributed in known weak enhancers and promoters.

### The gene regulatory network rewiring during senescence

Given that we have found some senescence specific chromatin regions, the accessibility of these regions gradually increased or decreased with senescence, and we have discovered their characteristics, then we need to answer the third question above that whether these regions are involved in the senescence fate decision. At first, we analyzed the genes with the most significant accessibility changes in these regions and displayed them in the form of heatmap (Fig. 3A). Among them, FOXA1, PLAT, CD44, FGF5, DIO2 and other genes are upregulated, whereas PLCB1, SETBP1, EHD3, ZNF423, PHGDH and other genes are downregulated in multiple senescence systems(Chan et al., 2019; Cho et al., 2019; Galanos et al., 2016; Hari et al., 2019; Hernandez-Segura et al., 2017; Komseli et al., 2018; Lau, Porciuncula, Yu, Iwakura, & David, 2019; Leveque et al., 2019; Q. Li et al., 2013; Limbad et al., 2020; Lowe & Raj, 2014; Nagano et al., 2016; Pan et al., 2019; Pantazi et al., 2019; Suwan et al., 2009; Wu, Pepowski, Takahashi, & Kron, 2019; Zhou et al., 2010), as are consistent with our results. These genes are located in or near the regions of IARs or DARs respectively. In order to explore the possible roles of IARs and DARs in senescence, we analyzed the signaling pathways that these genes are involved and the reported effects on individuals when their expression are perturbed. We found that IARs have a significant correlation with TGFβ pathway which suggested that IARs may participate in the regulation of TGFβ pathway and the most relevant physiological process is the morphogenesis of epithelial branch ducts. Nevertheless, the DARs have the strongest correlation with the metabolic process of proteoglycans and chondroitin sulfate. Further analysis for the disorder of related genes in mice showed that IARs perturbation may lead to pericardial effusion and epicardial morphological abnormalities while DARs perturbation may lead to aberration of kidney volume and morphological distortion of the fourth ventricle (Fig. 3B and Supplementary Fig. 6A). These comparisons indicated that the disorder of IARs and DARs related signaling pathways is not good for health. IARs are more closely related to senescence, and may regulate the senescence fate from the source in terms of the senescence related signaling pathways that involved.

**Fig. 3.**
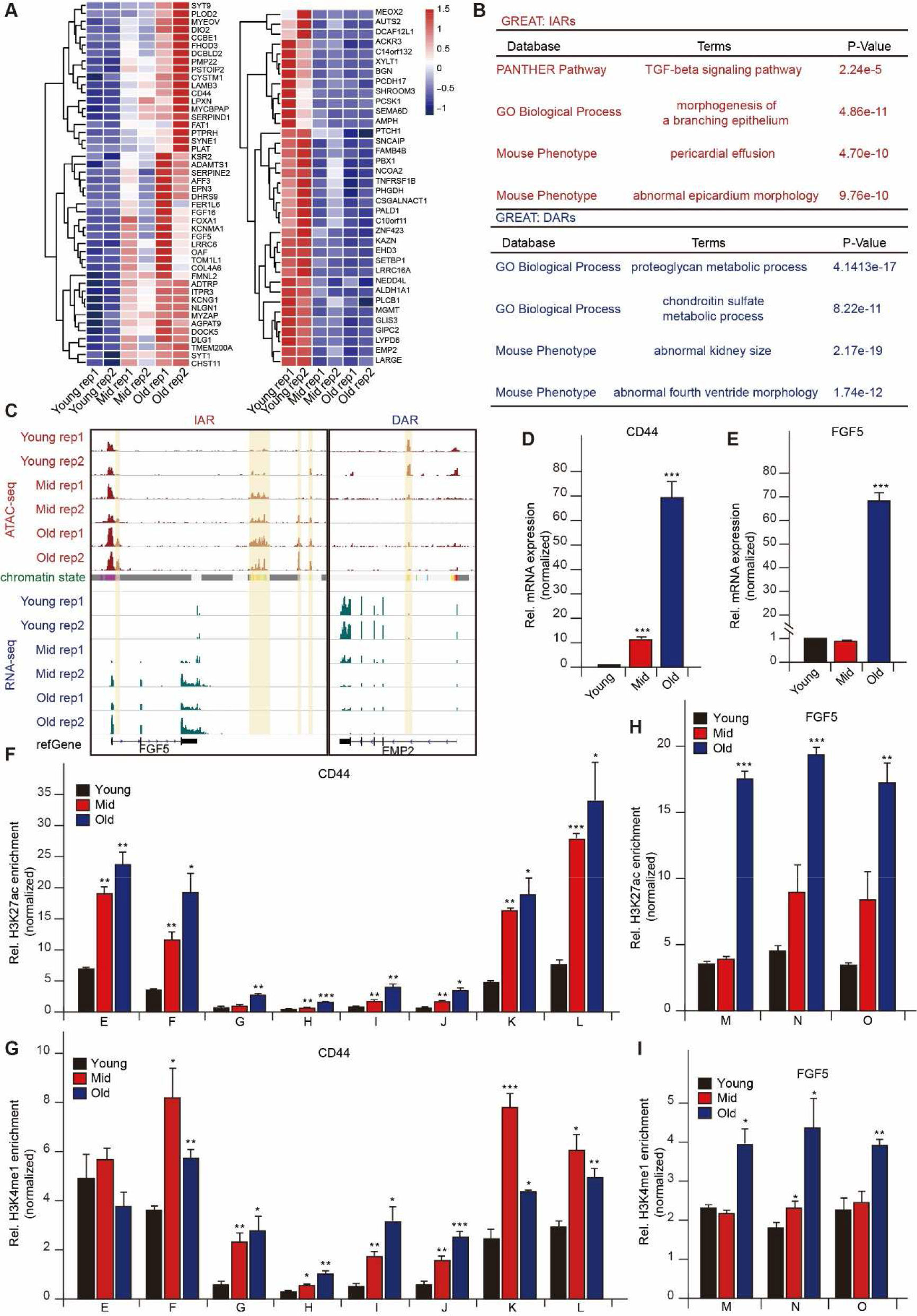
The gene regulatory network rewiring during senescence. **A** Heatmap showing the dynamic chromatin accessibility of remarkable genes belonging to IARs and DARs. **B** The analysis for GO enrichment and physiological process of IARs and DARs. **C** Snapshot showing the signal peaks of ATAC-seq and RNA-seq and adjacent genes near IARs and DARs. **D, E** Real-time qPCR results of CD44 (**D**) and FGF5 (**E**) showing the change of mRNA level during senescence. **F-I** Chromatin immunoprecipitation-qPCR (ChIP-qPCR) assays for IARs near or in the CD44 and FGF5. The dynamic change of H3K27ac enrichment of the IARs near CD44 (**F**) and FGF5 (**H**), and H3K4me1 enrichment of the IARs near CD44 (**G**), and FGF5 (**I**).

To show the regulation of IARs and DARs to related genes more vividly, we take FGF5, CD44, DIO2, SULF1, and EMP2 as examples to display the changes in ATAC-seq and RNA-seq signals near IARs and genes during senescence. There are 4 IARs regions near or inside FGF5 and the RNA-seq signals of FGF5 gradually increased with senescence (Fig. 3C). FGF5 is a classic secreted protein that inhibits hair elongation whose role in senescence remains unclear. However, FGF5 showed positive correlation with senescence in our results. CD44 is a typical SASP which mediates the recruitment of immune cells for of senescent cells in which two IARs are distributed and the CD44 expression also increased with senescence (Supplementary Fig. 6B). The same association existed in DIO2 (Supplementary Fig. 6C). Joana et al. also identified that DIO2 is highly expressed in the hMSCs senescence system, suggesting that DIO2 may be an alternative senescence marker(Medeiros Tavares Marques et al., 2017). Meantime, there is a DAR in EMP2 and its RNA-seq signals gradually weakened with senescence (Fig. 3C). EMP2 is a membrane protein responsible for important physiological functions such as cell adhesion and endocytosis which may function as the downstream effector of senescence. There are 7 DARs in or near the SULF1 and its expression also gradually decreased with senescence (Supplementary Fig. 6D). In addition, we aligned the H3K27ac signals and ATAC-seq signals of SULF1, found that the ATAC-seq signal peaks cover all H3K27ac signal peaks, but some regions that have no H3K27ac signals presented progressive ATAC-seq signals along with senescence development. These results demonstrated the importance of time dimension for measuring chromatin related characteristic signals, especially in progressive physiological processes such as senescence. More hidden information will be closed to the surface by comparing continuous senescence stages. It is worth noting that there distributes strong enhancer in the IARs near FGF5 (Buecker et al., 2014). This indicated that the IARs have a strong association with enhancers. The regulation of IARs for related genes may be mediated by the activation or silence of enhancers during senescence. Although some enhancers in IARs or DARs have been identified, there are still many enhancer-like regions have not been identified in which senescence related regulatory events possibly take place (Fig. 3C). In addition to paying attention to the common regulatory elements and chromatin accessibility features of IARs and DARs, we also studied the distribution pattern of DNA sequence on them. We found that the IARs has abundant repeated sequence which are mainly small RNAs, while DARs have few repeated sequence (Supplementary Fig. 6E). These results suggested that multidimensional regulatory events occur in IARs, including histones and DNA.

Then we found that IARs are evolutionarily conserved (Supplementary Fig. 7A), implying that these senescence specific regions may have stable functions which may contribute original driving force to senescence fate while DARs tend to correlate the function decline as a result of senescence. So, we focus our attention on the function and driver of IARs. We further used ChIP-qPCR and RT-qPCR to verify the accessibility of IARs and the expression level of adjacent genes during senescence. The information about relative position of the primers used in ChIP-qPCR in genome are displayed in Supplementary Fig. 7B. The mRNA levels of CD44 and FGF5 showed a characteristic of gradual increase with senescence (Fig. 3D and 3E). The signals of H3K27ac as a symbolic modification of chromatin accessibility enriched greatly in IARs of CD44 and FGF5 and presented a certain gradient (Fig. 3F and 3H). These results proved that our definition about IARs is credible, the chromatin accessibility of IARs did increase gradually along with the progresses of senescence. The enrichment of H3K4me1 as a marker of enhancer also increased remarkably with senescence in the IARs of CD44 and FGF5, but its increased mode is varied at different sites, which increased with continuous gradient at some sites while first increased and then decreased at other sites (Fig. 3G and 3I). Combining the analysis of H3K27ac and H3K4me1, our results indicated that the IARs has enhancer activity, but its activity is dynamically regulated during senescence. In summary, the reconstruction of chromatin accessibility predetermines the gene expression program during senescence, there are senescence specific regions IARs and DARs whose accessibility are regularly changed. In particular, IARs are closely related to the occurrence of senescence, the regulation of IARs to senescence mediated by special elements with enhancer activity in IARs.

### DNA methylation may contribute to establish chromatin accessibility

Considering that the overall accessibility of chromatin undergoes regular rearrangement during senescence, this is very similar to the reconstructed pattern of DNA methylation during senescence. The DNA methylation of whole genome presents a broad decline and a local increase(H. A. Cruickshanks et al., 2013). So, we wondered investigate whether there is connection between DNA methylation and chromatin accessibility during senescence, and the regulatory relationships between them, and whether the IARs and DARs have a correlation with DNA methylation during senescence. First, we extracted the DNA methylation data (GSE82234) in HUVECs senescence from database(Franzen et al., 2017) to align our ATAC-seq data, we found that there is an obvious negative correlation between the DNA methylation signals and ATAC-seq signals during senescence. When DNA methylation increased, chromatin accessibility decreased, and vice versa (Fig. 4A). Then we analyzed the distribution pattern of DNA methylation in genome among different senescence periods, and found that the distribution of DNA methylation in different periods is almost same. The DNA methylation signals near TSS is very low, but they are very high far away TSS (Fig. 4B). Its distribution regularity has a strong correspondence with ATAC-seq signals, the enrichment of ATAC-seq signals near TSS is the highest that is much higher than the area far from TSS (Supplementary Fig. 4B). At the same time, the chromatin accessibility and DNA methylation are also opposite to each other at specific sites, such as inside or near DNAJC8, ATPIF1, COL16A1, the ATAC-seq signals is very low at the site where the DNA methylation signals is high and vice versa (Fig. 4C). Such an obvious trade-off relationship strongly suggested that there may be causality between DNA methylation and accessibility. To further explore whether the dynamic changes between DNA methylation and chromatin accessibility during senescence are consistent, we analyzed the changes of DNA methylation in IARs and DARs whose accessibility regularly increased or decreased during senescence. We found that the DNA methylation level in the IARs was significantly lower than that in the DARs (Fig. 4D). For instances, the ATAC-seq signals gradually increased while the DNA methylation signals gradually decreased in IARs of CD44 along with senescence. On the contrary, The ATAC-seq signals gradually decreased while the DNA methylation signals gradually increased in DAR of EMP2 along with senescence (Fig. 4E). This indicated that there is a potential regulatory relationship between chromatin accessibility and DNA methylation during senescence. Researches have shown that DNA methylation can induce the chromatin condensation, especially in the process of replicated senescence. Both the DNA methylation level and the condensed chromatin proportion were significantly decrease(Choy et al., 2010). In summary, our results indicated that the reestablishment of DNA methylation patterns are involved in the chromatin accessibility remodeling, and that changes in DNA methylation are more likely to occur before chromatin accessibility. The decrease of DNA methylation contributes to the enhancement of chromatin accessibility at specific sites.

**Fig. 4.**
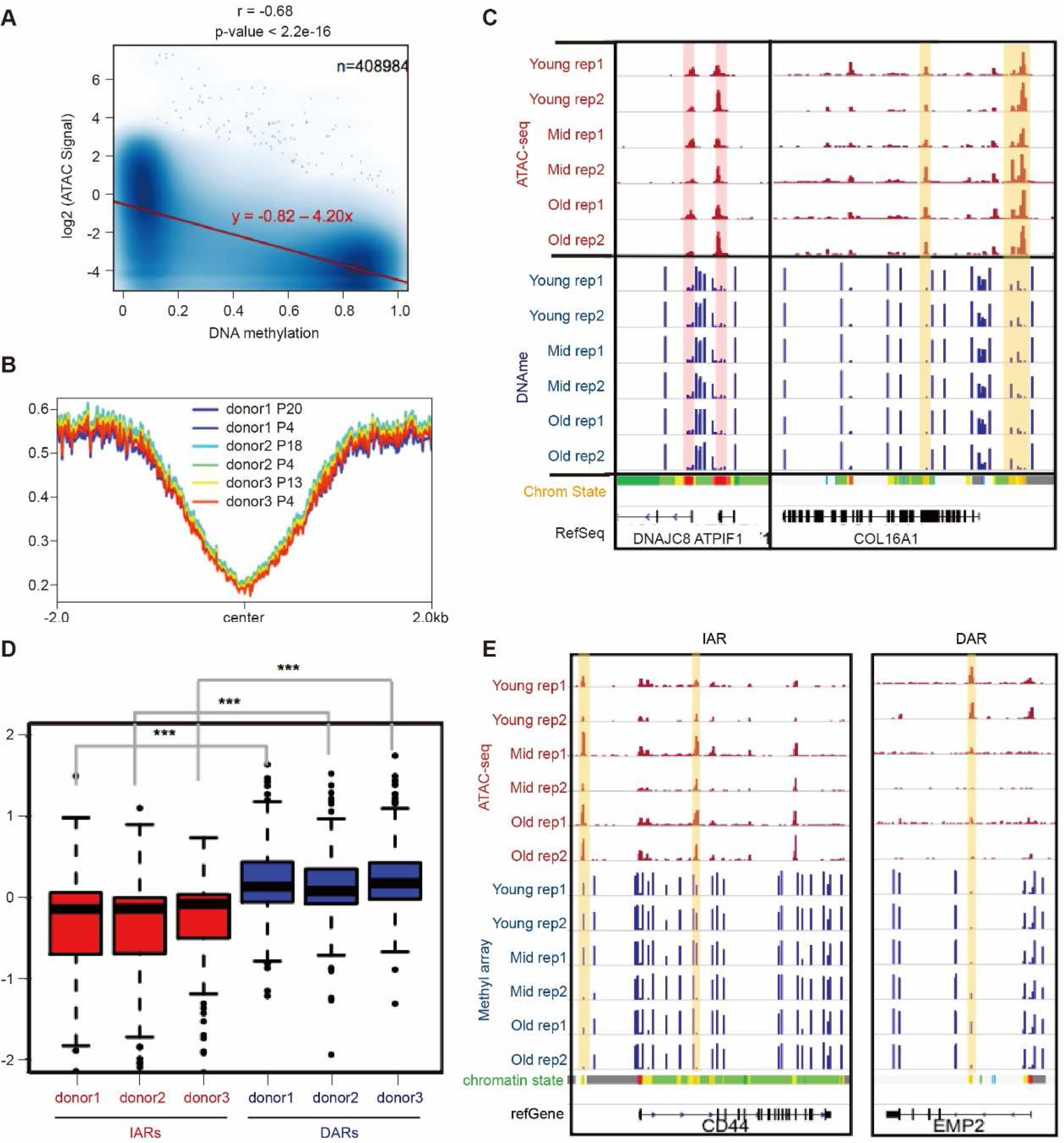
DNA methylation may contribute to establish chromatin accessibility. **A** The correlation analysis for the chromatin accessibility and DNA methylation in HUVECs senescence. **B** The distribution of DNA methylation signals in different senescence stage of HUVECs. **C** Snapshot showing the correspondence between ATAC-seq signals and DNA methylation signals near or in the DNAJC8, ATPIF1 and COL16A1. **D** The box plots showing the DNA methylation level in DARs and DARs. **E** Snapshot of IAR and DAR examples showing the change patterns of the DNA methylation and ATAC-seq signals near or in the CD44 and EMP2.

### AP-1 is a potential regulator to regulate the accessibility of IARs in HUVECs senescence

Given that chromatin accessibility remodeling predetermines the gene expression profile in senescence, we were sought to explore the key factors that drive the chromatin remodeling, which may help us uncover the senescence secrets. Some special transcription factors often play a leading role in chromatin structure remodeling. To investigate which transcription factors may be involved in the rearrangement of chromatin accessibility during senescence, we performed the motif enrichment analysis for transcription factors in IARs and DARs through the two algorithms HOMER and MEME. Surprisingly, the enrichment of AP-1 family was much more significant than other transcription factors in IARs. These AP-1 transcription factors are mainly ATF3, FOSL1, FOSL2, BATF, and JUNB, among them, the enrichment of ATF3 was the highest (Supplementary Fig. 9D). Compared with the remarkable enrichment of transcription factors in IARs, that in DARs are less significant where ETV1, ETS, GABPA, ERG, FLI1 are mainly distributed (Fig. 5A, Supplementary Fig. 9A and 9B). Combined with the classic senescence signaling pathways that may be regulated by IARs, such as the TGFβ pathway (Fig. 3B), we believe that the AP-1 family is the driver of chromatin accessibility remodeling during senescence, and their reconstruction for the chromatin regions is selective. At the same time, in order to verify the conclusion that DNA methylation negatively regulate the chromatin accessibility during senescence, we analyzed the methylation level in major motif sites of AP1 transcription factor. We found that the methylation level in open region is lower than closed region (Fig. 5B), which indicated that DNA methylation affect the accessibility reconstruction of neighboring chromatin by controlling the binding of transcription factors to their motifs.

**Fig. 5.**
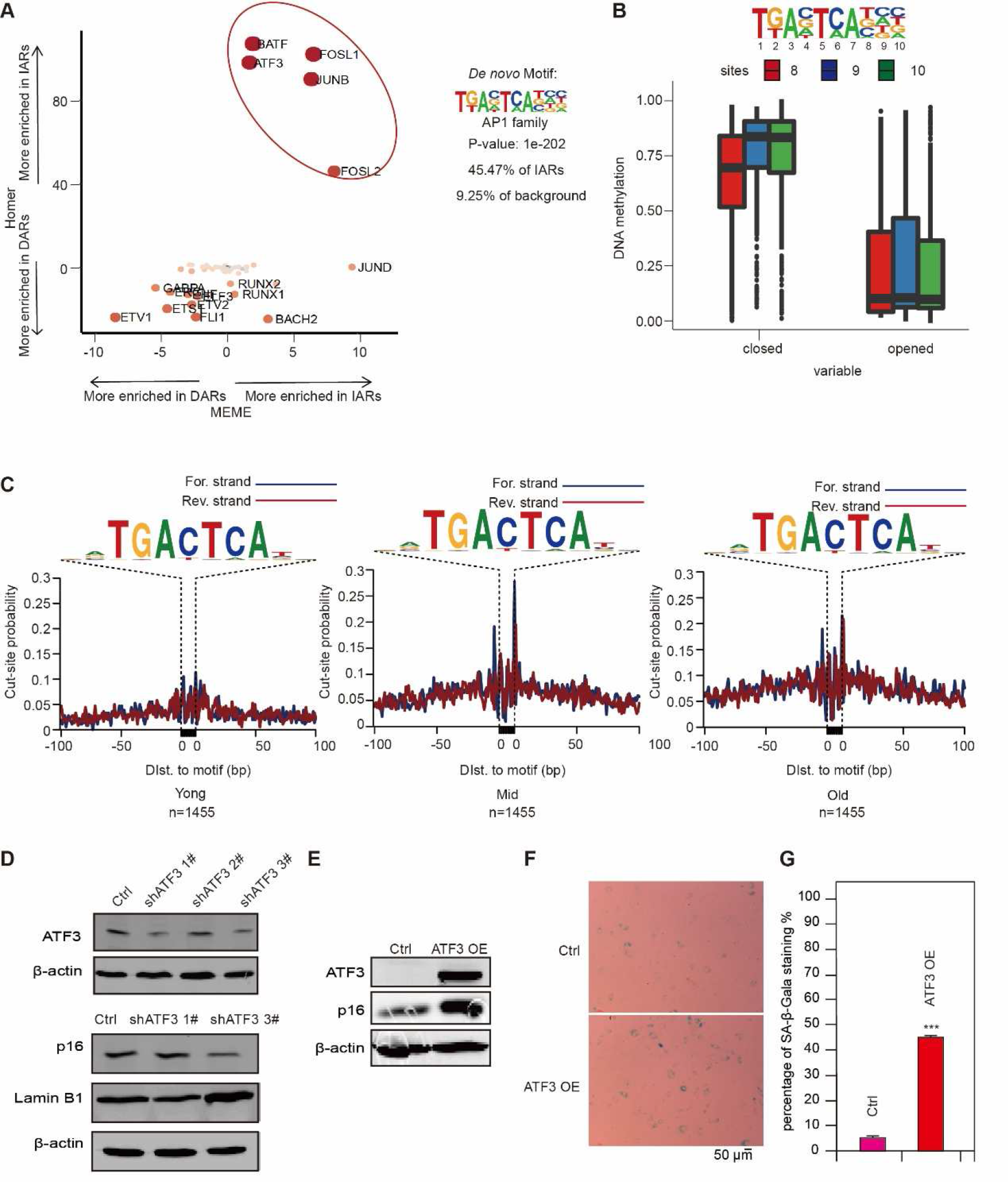
AP-1 is a potential regulator to regulate the accessibility of IARs in HUVECs senescence. **A** Motifs analysis for open chromatin in HUVECs senescence with HOMER and MEME algorithms. **B** Box plot showing the difference of DNA methylation at 8, 9, 10 sites of the AP-1motifs between closed and open chromatin. **C** The ATAC-seq footprints analysis for ATF3 along with the senescence of HUVECs. **D** Western blot showing the protein levels of ATF3, p16, Lamin B 1 after knocking down ATF3 in HUVECs. β-actin served as loading control. **E** The protein expression of ATF3 and p16 after ATF3 overexpression in HUVECs. β-actin served as loading control. **F** SA-β-Gal staining assay showing the difference of senescence degree between ATF3-overexpressed and controlled HUVECs. **G** The diagram of statistics for SA-β-Gal positive cells in (**F**).

Among the AP-1 transcription factors enriched in IARs, the highest ranked ATF3 attracted our attention. Next, we want to explore the role of ATF3 in senescence fate decision and whether it is a desired target for intervention to senescence. We found that the ATAC-seq signals at the ATF3 binding sites gradually increased along with the progress of senescence (Fig. 5C). In combination with the high enrichment of ATF3 in IARs, our results indicated that the ATF3 significantly improve chromatin accessibility near its binding site during senescence. Then we investigated whether ATF3 promotes the occurrence of senescence, and whether ATF3 defect delays senescence or reduce the expression of senescence related markers. We performed the knockdown and overexpression experiments of ATF3 in HUVECs. We found that the protein level of P16, a marker of senescence, was downregulated, and the protein level of LMNB1 was upregulated after ATF3 defect. Conversely, when we overexpressed ATF3, the P16 protein level is upregulated and the SA-β-Gal signal is enhanced, which indicated that ATF3 can indeed promote senescence. Taken together, the AP-1 family, especially ATF3, is involved in the establishment of senescence program, most likely by regulating the accessibility of enhancers or other regulatory elements in IARs thereby reconstructing the senescence related gene expression profile.

## Discussion

Our study indicated that the gene expression profile in senescent cells is predetermined by accessible chromatin rearrangement. The senescence-specific chromatin regions IARs and DARs have distinct functions for senescence. The IARs are involved in the regulation for senescence which are mainly distributed in defined heterochromatin regions. The DARs are related to the decline of cell function caused by senescence which contain weak enhancers and promoters. More importantly, we identified that the gradual increase in accessibility of IARs during senescence is primarily driven by AP-1 transcription factors, especially the ATF3. The reshaping of chromatin accessibility by AP-1 transcription factors such as ATF3 is affected by DNA methylation at their binding sites. In general, ATF3 has a significant effect in the reconstruction of gene expression program and further promoting senescence, which is mediated by regulatory elements such as enhancers in IARs. Therefore, ATF3 is at the top of the senescence regulatory network and may be an effective target for senescence intervention.

The senescence fate depends on epigenetic pattern, as is a systemic state in where chromatin remodeling, histone modifications and DNA methylation cooperatively decide the way ahead of cells. But the understanding about their interaction and priority is still fragmentary. We systematically analyzed the chromatin accessibility remodeling in different stages of senescence using ATAC-seq technology. Our results demonstrated that senescence complies a slated program, as is mainly scheduled by chromatin architecture. The consistent trajectory between chromatin accessibility and gene expression during senescence shows a possible regulatory direction that senescence-specific gene expression profile is determined by chromatin rearrangement pattern (Fig. 1F and Fig. 3A). For a long time, the panorama of senescence remains to be elucidative. It increasingly seems that chromatin architecture remodeling may be a united theory to describe a series of events in senescence. Based on our results about the sequential and continuous increase of chromatin accessibility and gene expression during senescence (Fig. 3), we proposed that senescence executes a cascade program comprising preparation, initiation, promotion and deterioration. In various stages of senescence, perhaps the respective gene expression patterns are built by combining different regulators. It’s necessary to take correspondent measures for different senescence stage considering the dynamic but traceable epigenetic landscape and gene expression profile in senescence. As a holistic process, senescence involves multiple pathway related to plenty of genes. Changing the expression of multiple senescence related genes at the same time may be an effected strategy for senescence intervention. A. Ocampo et.al ameliorated senescence and aging phenotype by short-term cyclic expression of Oct4, Sox2, Klf4, and c-Myc (OSKM)(Ocampo et al., 2016). T. J. Sarkar et.al transiently expressed OCT4, SOX2, KLF4, c-MYC, LIN28, and NANOG (OSKMLN) in human cell partially reverted cellular senescence and restored aged tissue(Sarkar et al., 2020). These factors withdraw the senescence program by bringing broad changes to chromatin landscape that further rearranging the gene expression profile. Our whole-genome chromatin accessibility and gene expression landscape provides global perspective to senescence which could generates general orientation to find intervention target for senescence and aging. In our research, chromatin accessibility is fluctuant away from TSS and most of them lie in intronic and intergenic regions in where distal regulatory elements are plentiful (Fig. 1G, Fig. 3C-I, Supplementary Fig. 5D-E and Supplementary Fig. 6B-D). It’s suggested that many senescence associated regulatory elements may be hidden in distal intergenic regions rather than proximal regions.

Above all, we identified the senescence-specific chromatin regions IARs and DARs. In particular, IARs have conspicuous characteristics related to the occurrence of senescence, and we found that the drivers facilitating these special regions are AP-1 transcription factors which function as pioneer-like transcription factors. This is a novel role for AP-1 family and rare researches report the relationship between AP-1 and senescence. Meanwhile, we noticed that the heterochromatin contributes most of IARs (Fig. 2E) which shows a possibility that the disappeared heterochromatin was transformed to open chromatin through gaining the accessibility, indicating the potential special contribution of heterochromatin for senescence. As is consistent with heterochromatin loss model in which heterochromatin disassociate from lamina (LAD). Meantime, the decrease of DNA methylation level at the AP-1 binding sites promotes its enrichment in IARs, which successfully reshapes the accessibility pattern of adjacent chromatin (Fig. 4C-E and Fig. 6B). Our results provide a novel clue to the relationship among the heterochromatin, DNA methylation and senescence. When our project was about to be completed, we found that Ricardo et al.(Martinez-Zamudio et al., 2020) recently published research on the regulation of AP-1 transcription factors to senescence program and our work can confirm and complement each other. The senescence system they chose is the oncogene induced senescence (OIS) in WI-38 while we chose the replicated senescence of HUVECs. Although using different cell lines and different types of senescence, we both found that the AP-1 transcription factors family can regulate the senescence fate by remodeling the gene expression profile, and perturbing the expression of AP-1 transcription factors can reverse the senescence program. However, they identified c-JUN as the core AP-1 transcription factor that regulates senescence different from the ATF3 in our research. This shows that the AP-1 transcription factors play a role as a driver in senescence, but in which different AP-1 transcription factors are involved and there may be different combination modes between them in different cell types and senescence types. It is necessary to conduct more exploration for the functions of AP-1 family in senescence to discover more specific targets and intervention strategies for aging. The chromatin accessibility of IARs changed regularly during senescence where a large number of known enhancers and the regions that have enhancer activities but not been identified are distributed. And the enrichment of AP-1 transcription factors such as ATF3 in IARs gradually increased with senescence. Thus, we proposed that AP-1 transcription factors remodel the gene expression program through regulating the accessibility of enhancers and other regulatory elements in IARs, and finally leads to senescence. It’ a coincidence that the work of Ricardo et al. showed that the activity of a batch of enhancers gradually increased during senescence, which are consistent with our conclusion that the senescence program is preset in cells and the senescence fate is predetermined by specific gene expression program that is reconstructed by some key upstream transcription factors through regulating the activities of special enhancers or other regulatory elements.

There were recent studies implied that AP-1 could drive breast cancer and oncogene induced senescence by regulating accessibility of specific promoter and enhancer(Han et al., 2018; Vierbuchen et al., 2017). Functional CRISPR screen identifies that AP-1 associated enhancer modulates oncogene-induced senescence by regulating FOXF1^97^. Combing previous researches and our results, we proposed that AP-1 is a significant pro-senescence regulator. There are still some detailed mechanisms to be pored about the regulation of chromatin accessibility redistribution by AP-1 during senescence, such as interaction network and action time. And more chromatin landscape in multiple senescence systems are needed.

After all, we presented a sequentially dynamic landscape orchestrating the chromatin accessibility and gene expression during entire culture process for human primary cells *in vitro*. In which we defined IARs and DARs that are two kinds of senescence-specific chromatin regions. DNA methylation possibly promotes the reopening of heterochromatin which contribute IARs for senescent cells. DNA methylation affect the binding of AP-1 transcription factors to the regulatory elements in IARs. We described that the AP-1 transcription factors, especially ATF3 promotes senescence by remodeling the chromatin accessibility pattern in IARs. Accordingly, our research provides a potential target to senescence and aging.

## Methods

### Cell culture

Primary human umbilical vein endothelial cells (HUVECs) were purchased from ALLCELLS Co. Ltd., Shanghai, China and cultured in endothelial cell medium (ECM, Sciencell, USA). HEK293T cells purchased from American Type Culture Collection (ATCC) were cultured in DMEM (GIBCO, Grand Island, NY, USA), supplemented with 10% fetal bovine serum (FBS, GIBCO, Grand Island, NY, USA). All cells were routinely maintained at 37 °C in a humidified incubator with 5% CO2. HUVECs were passaged at 80% to 90% density in 10 cm dish coated with 0.25% gelatin that usually need 2 or 3 days with 0.5×10^6^ start-up cells. HUVECs were cultured starting from passage 2 (about PD4) until they didn’t proliferate anymore.

### ATAC-seq

Freshly cultured cells at different time point of the senescence time course were collected. Cell pellet was resuspended with moderately cold lysis buffer (eg, 1.0×10^^6^ cells, add 400 μl lysis buffer). 7.0×10^^4^ cells were transferred to sterile PCR tube for subsequent transposition reaction and library construction using TruePrep DNA Library Prep Kit V2 (#TD501, Vazyme Biotech Co., Ltd, Nanjing, CHN). 50μl transposition mixture containing 5× TTBL and TTE mix V50 (TD501) was slowly dripped into the cell suspension and mix thoroughly together, then incubated that for 30 min at 37°C to perform the DNA fragmentation and adaptor addition. DNA was purified using Qiagen MinElute PCR purification Kit (#28004, Qiagen, Germany) followed the handbook. DNA precipitation was dissolved in 26 μl ddH2O as the template for amplification. DNA libraries were produced by PCR amplification 8 cycles using the above library preparation kit. The products were purified and DNA precipitation was dissolved in 10 μl 10 mM Tris HCl, pH 8.0. Paired-end sequencing was performed in an Illumina Hiseq PE150 (Novogene, Beijing, CHN).

### RNA-seq

Total RNA from different passaged cells were extracted by TRIzol reagent (#15596018, Invitrogen, USA). RNA library construction and Illumina sequencing with paired-end Hiseq PE150 were performed in Novogene Biotech Co. Ltd., Beijing, China.

### Senescence-associated β-galactosidase (SA-β-gal) staining

Before the SA-β-gal staining assay (Sigma, CS0030, USA), about 0.6×10^^5^ cells were seeded in 6-well plates. Briefly, the adherent cells were washed twice with 1 ×PBS after aspirating the media. Cells were fixed with 1×Fixation Buffer and incubated for 7 minutes at room temperature. Then prepared staining mixture was added to all experimental wells after rinsing the cells 3 times with 1 ×PBS. Next, the fixed cells with staining mixture were incubated overnight at 37°C. The images were acquired using inverted microscope with 10 × 10 magnification (Leica DMI 6000B, Leica, Germany). Image J software (NIH) was used to analyze the SA-β-Gal signals.

### Immunofluorescence

About 0.5×10^^6^ cells were seeded in 10 cm dishes with sterile cover slides and cultured about 48h. The cells on cover slides were fixed for 10 min at room temperature with 4% paraformaldehyde after washing twice with cold 1× PBS. Then the cells were blocked for 30 min with 1% BSA at room temperature. After washing with 1× PBS, the cells were incubated overnight at 4 °C with 80 μL primary antibodies diluted in PBST (anti-Ki67, dilute at 1:500, ab15580, Abcam). Later, the cells were washed 4 times with 0.1% PBST and incubated for 2 h at room temperature with 80 μL secondary antibodies diluted in PBST (Alexa Fluor 594 Donkey Anti-Rabbit, dilute at 1:500, Life Technologies). After washing with PBST, the cells were incubated for 3 min with 1 ng/μL DAPI at room temperature. Last, the cover slides were sealed with 10 μL Fluoromount-G. One hour later, images were aquired using fluorescent microscope. Image J software (NIH) was use to analyze the immunofluorescence signals.

### Chromatin immunoprecipitation (ChIP)

The cells of for ChIP assay were cultured till about 80% confluence. 1% formaldehyde and 0.125 M glycine were used to initiate or terminate crosslinking, seperately. The scraped cells were collected to 1.5 mL DNA LoBind Tube (# 022431021, Eppendorf). Centrifuge the samples for 5 min at 4 °C with 2500 g and resuspended pellete with nuclei lysis buffer (50 mM Tris-Cl, 10 mM EDTA, 1%SDS and protease inhibitors cocktail). Then, the released chromatin was sonicated using bioruptor (Diagenode, Belgium) and diluted with IP dilution buffer (20 mM Tris-Cl, 2 mM EDTA, 150 mM NaCl, 1% Triton-X100 and protease inhibitors cocktail). The samples were incubated with specific antibodies overnight at 4°C. The antibodies were anti-H3K4me1(ab8895, Abcam), anti-H3K27ac (ab4729, Abcam), anti-H3 (ab1791, Abcam). The chromatin-antibody complexes were captured by protein A / G beads. qPCR assay was used to quatified the immunoprecipitated DNA. The primer information used in this analysis are included in the Supplementary Table.

### Real time and quantitative PCR (RT-qPCR)

Total RNA was extracted from cultured cells at 80% confluence using TRIzol Reagent (#15596018, Invitrogen, USA) according to the instruction. Then the RNA resolved in DEPC-treated water was reverse-transcribed to cDNA using All-in-One Supermix (#AT341, TransGen Biotech Co. LTD., Beijing, CHN). The cDNA levels were analyzed by qPCR (LightCycler, Roche, Swiss) with specific primers. The results were normalized to β-actin. All qPCR related primers are available in Supplementary Table.

### Lentiviral production and viral transduction

The ATF3 shRNA constructs were purchased from Sigma. The pSIN-3×flag-ATF3 vector was derived from pSin-EF2-Nanog-Pur (#16578, Addgene). HEK293T cells were seeded so that cells density could be 80% confluence when it comes to transiently transfection. The target construcs, psPAX and pMD2.G were co-transfected using calcium phosphate. The media were replaced with DMEM supplemented with 30% complement inactivated FBS after 18h transfection. The viral supernatants were collected 30h later. HUVECs were infected by the packaged viruses with 5 μg/mL polybrene (#MC032, M&C Gene Technology, CHN) when cell density come to 60% confluence. 36-48 h after infection, transducted cells were selected with 0.5 μg/mL puromycin (MA009, M&C Gene Technology, CHN)..

### Western blot

Proteins were extracted by TRIzol Reagent (#15596018, Invitrogen, USA) and resolved in 1% SDS. After the SDS-PAGE, the samples were transferred to nitrocellulose membranes and incubated for 30min at room temperature with 5% skimmed milk. The membranes were incubated overnight with specific primary antibodies at 4 °C. The primary antibodies were anti-LMNB1 (1:1000, 12987-1-AP, proteintech), anti-β-actin (1:2000, sc-47778, Santa Cruz), anti-p16 (1:500, ab108349, Abcam), anti-ATF3 (1:500, ab207434, Abcam). The membranes were incubated 2h with secondary antibodies IRDye800CW Goat/Donkey anti-Mouse/Rabbit (1:10000, 926-32210, LI-COR,) at room temperture after washing 3 times with PBST (1× PBS add 0.1% Tween-20). The blot information was aquired by Odyssey Infrared Imaging System (Odyssey, LI-COR).

---

**Supplementary Fig. 1.**
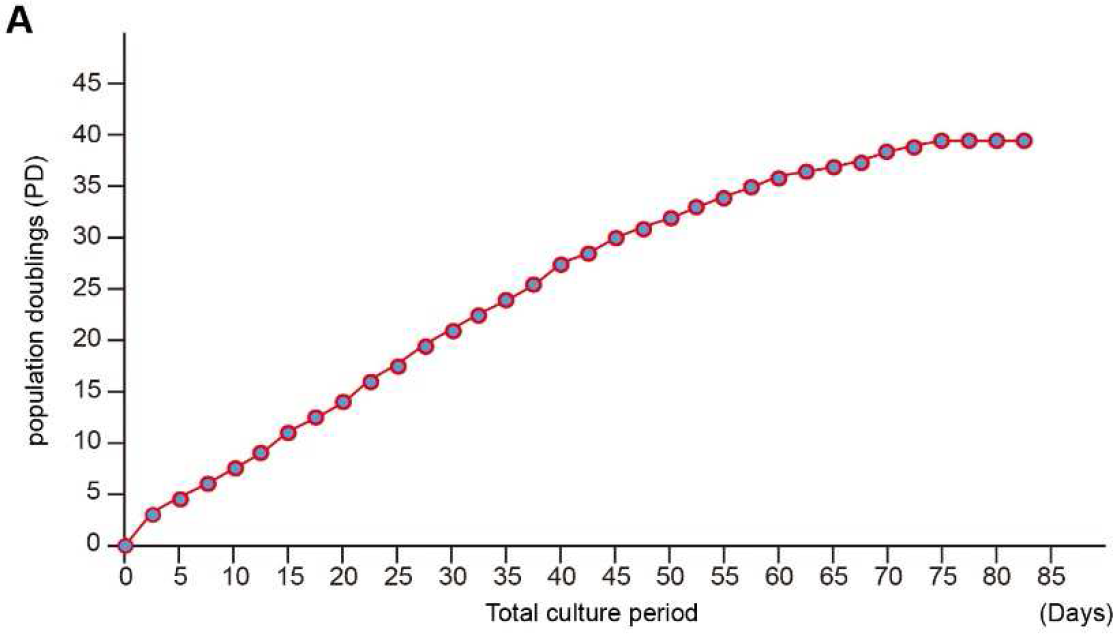
**A** The growth curve of HVUECs cultured *in vitro*. The HUVECs were cultured according to the inoculation amount of 0.5×10^6^ cells and grow to 80% to 90% confluence for one passage, and count the number of cells. The calculation formula of PD is: PD=log(N/N0)+S, where N represents the amount of inoculation in a specific period, N0 represents the initial amount of inoculation, and S represents the initial cell proliferation generation.

**Supplementary Fig. 2.**
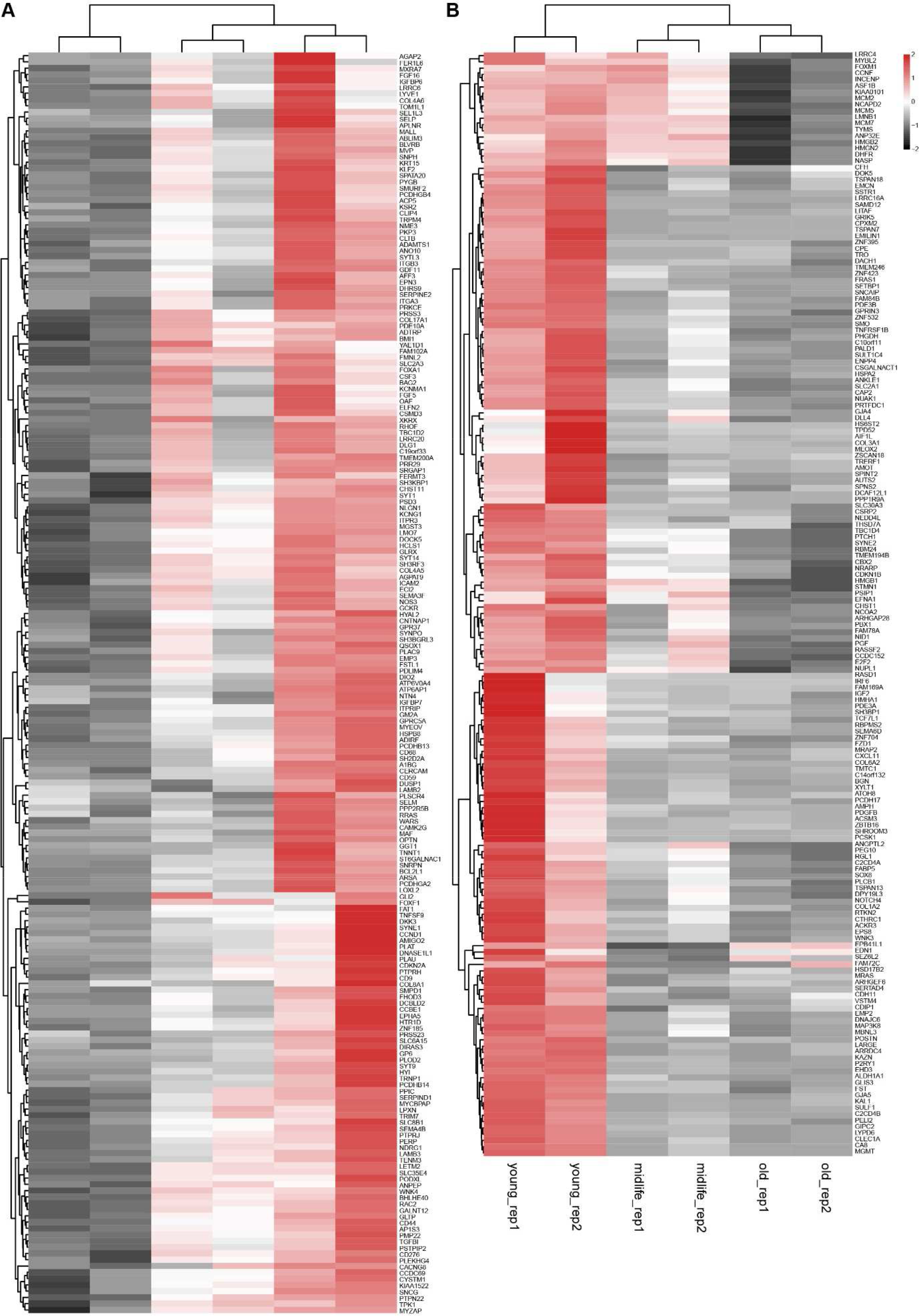
The heatmaps of RNA-seq showing the differential genes in HUVECs senescence. **A** The significantly upregulated genes during senescence. **B** The significantly downregulated genes during senescence.

**Supplementary Fig. 3.**
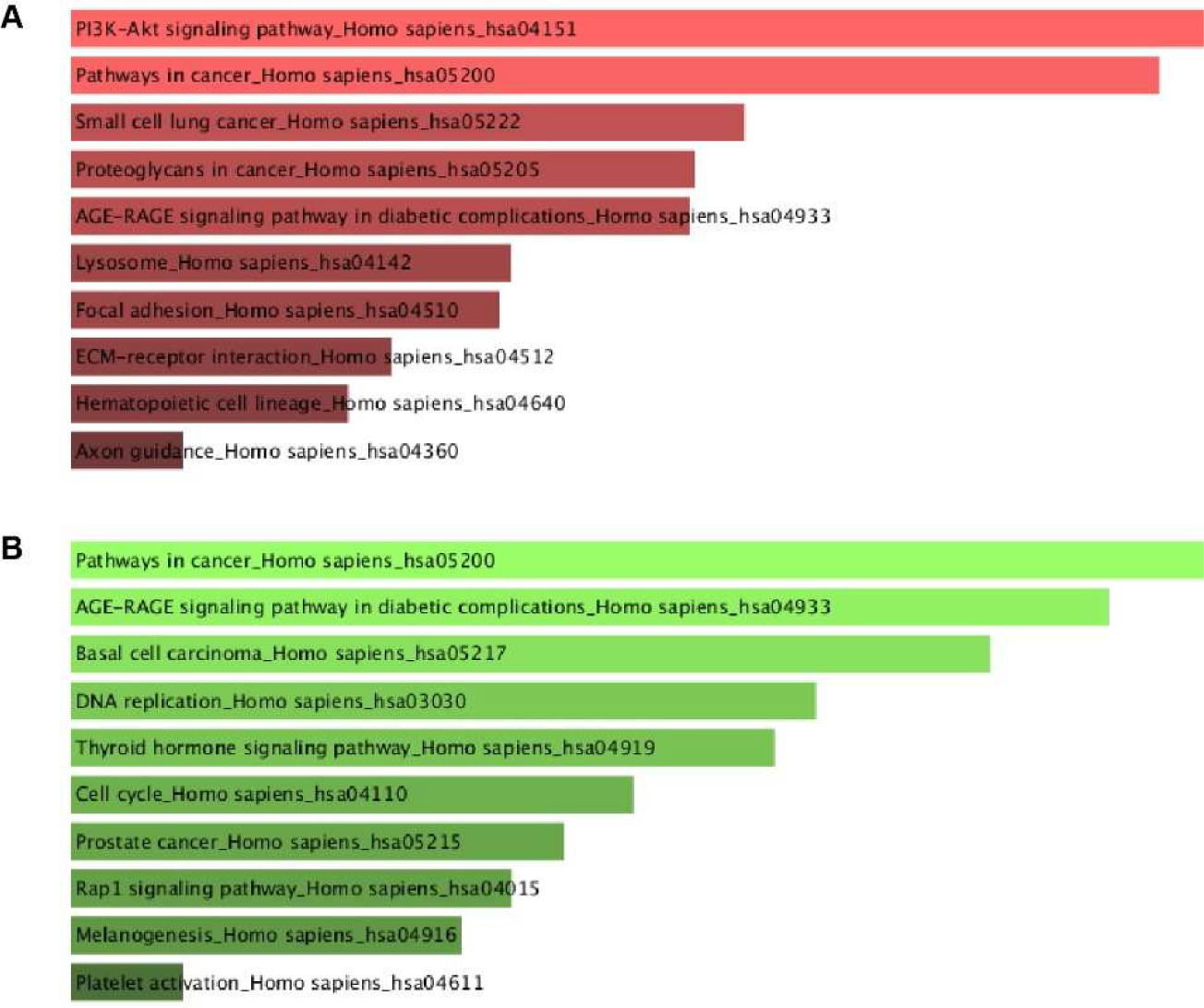
The GO analysis for differential genes in HVUECs senescence. **A** The upregulated signaling pathways during senescence. **B** The downregulated signaling pathways during senescence.

**Supplementary Fig. 4.**
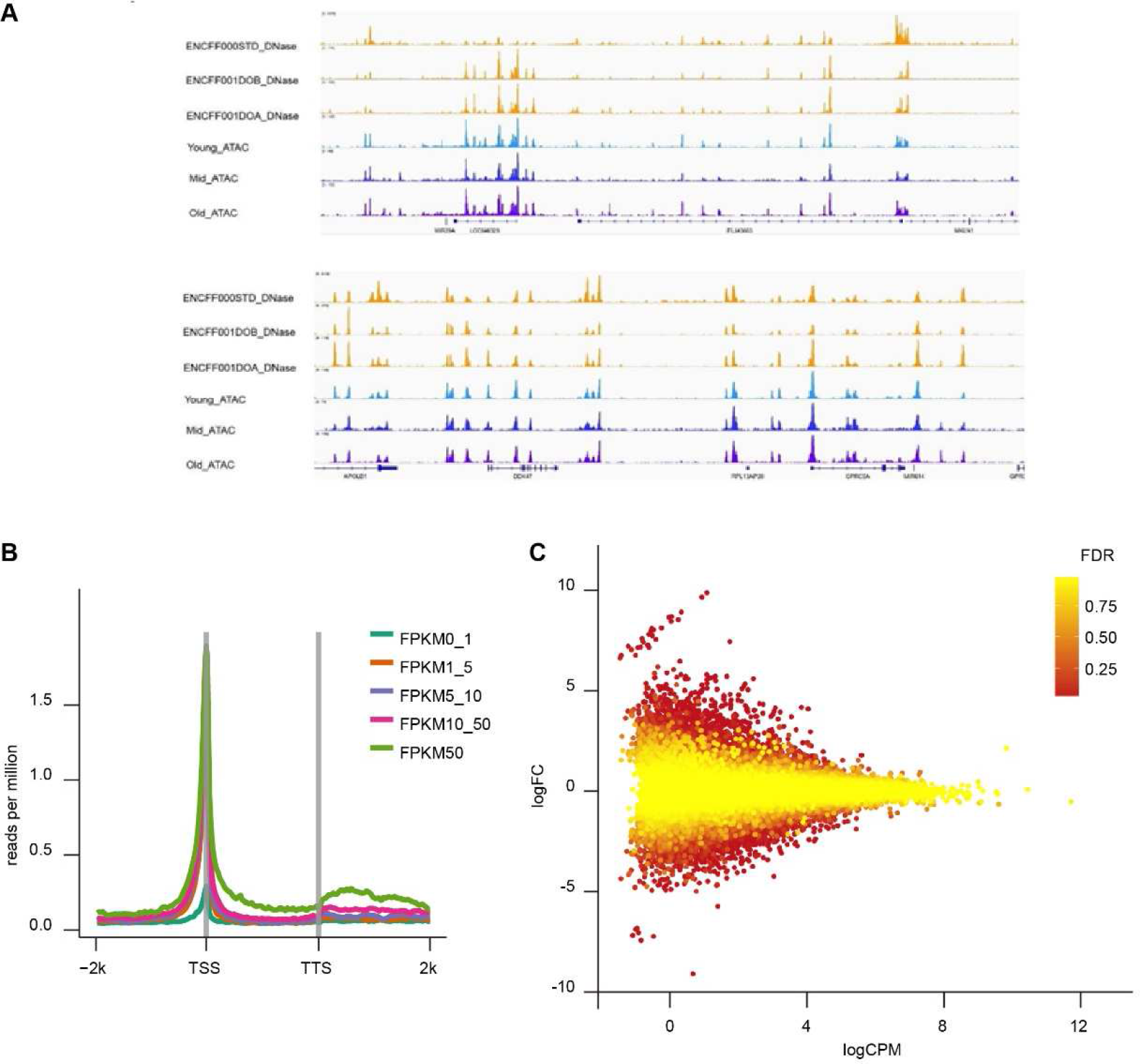
The analysis for the quality of ATAC-seq data. **A** The alignment between ATAC-seq and DNase-seq. **B** The analysis for the FPKM of ATAC-seq signals. **C** The scatter plot o ATAC-seq signals.

**Supplementary Fig. 5.**
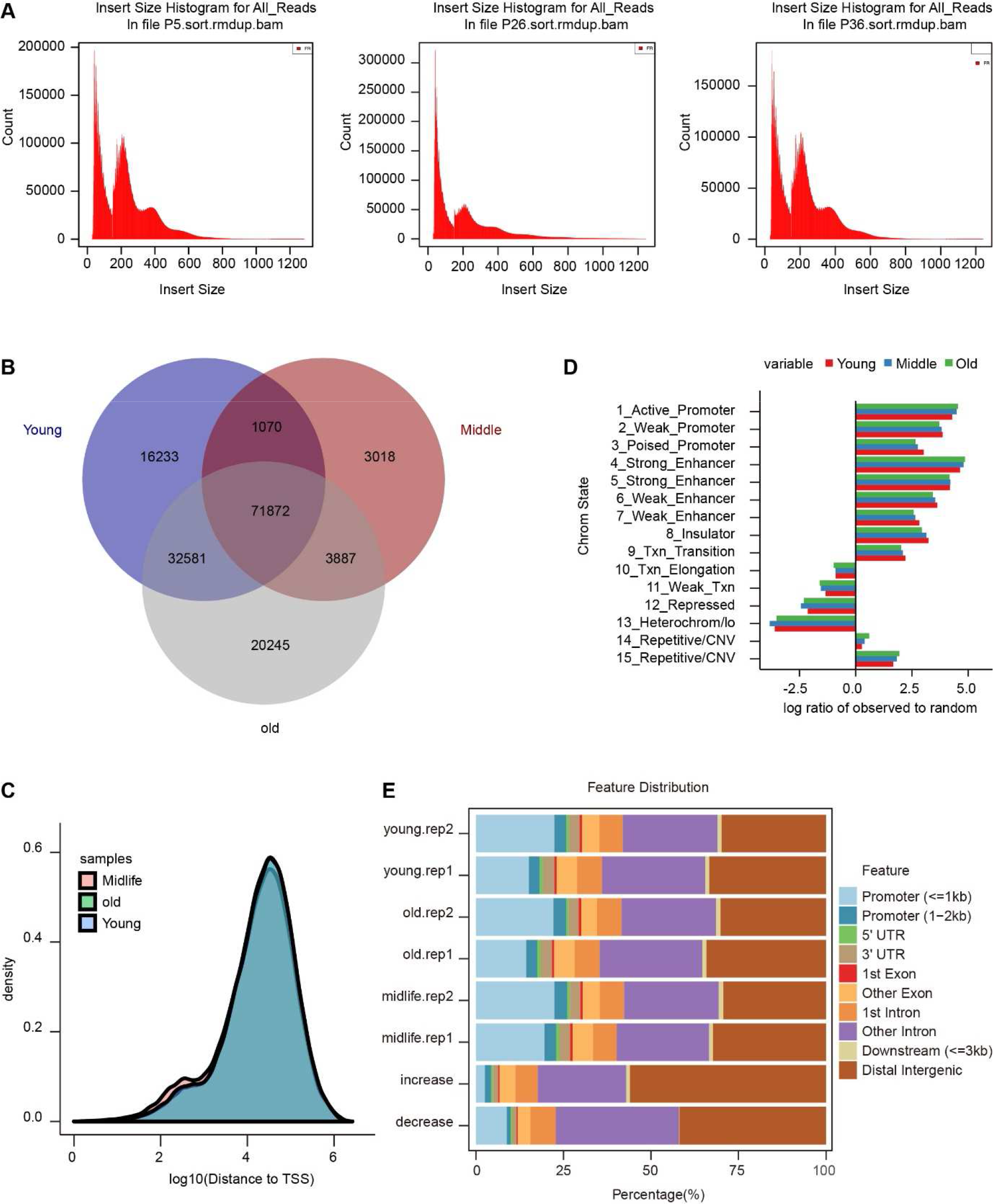
The characteristics of the distribution of ATAC-seq signals among different senescence stages. **A** The statistics for the ATAC-seq signal numbers in young, middle and old HUVECs. **B** The venn plot showing the overlap of ATAC-seq signals among three senescence stages. **C** The density plot of ATAC-seq signals showing the distribution rule distance to TSS. **D** The statistics of chromatin states for significantly changed regions of chromatin accessibility during senescence. **E** The feature distribution of the changed accessible chromatin regions during senescence.

**Supplementary Fig. 6.**
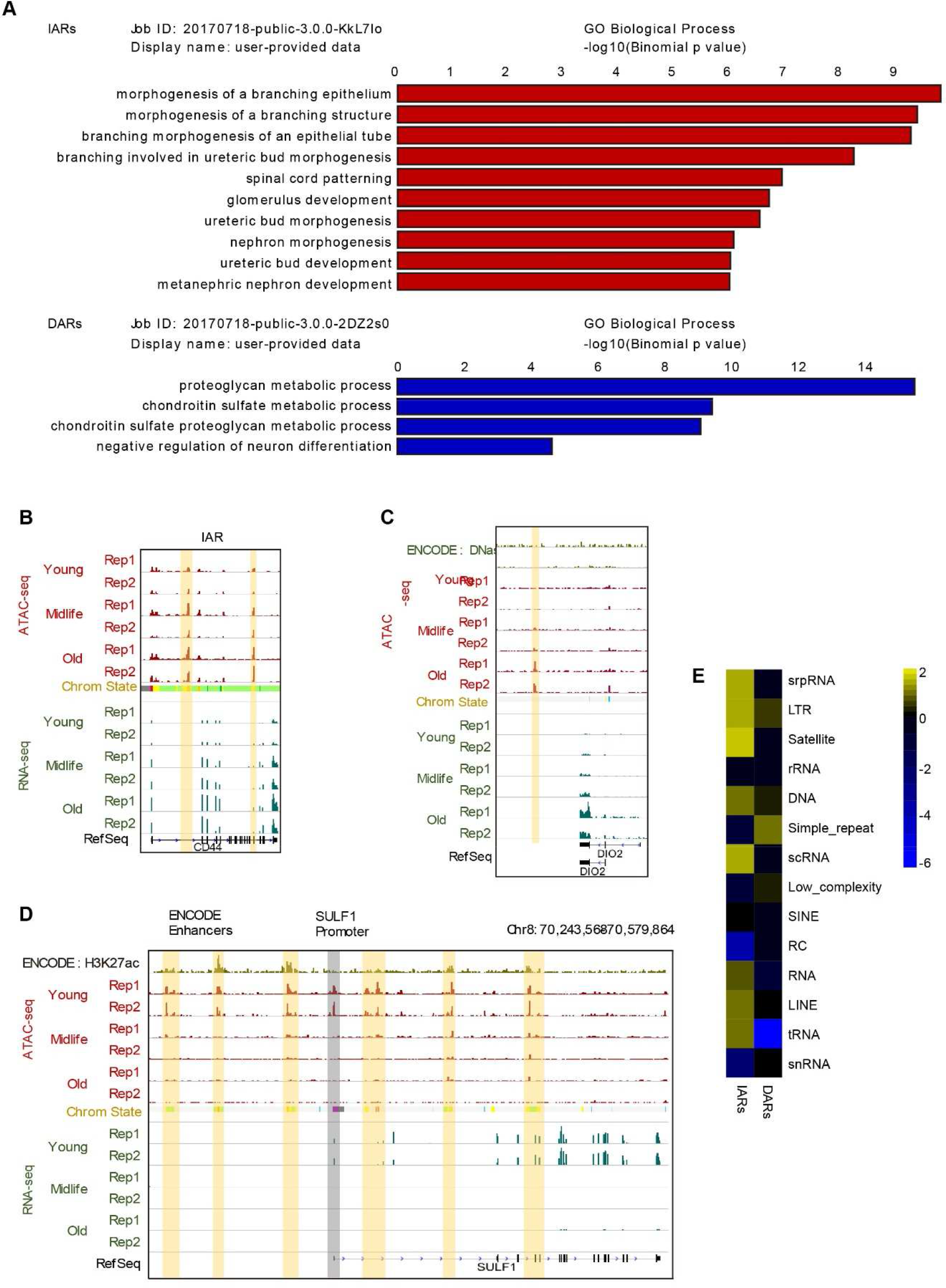
The analysis for the characteristics of IARs and DARs. **A** The GO analysis for the IARs and DARs related genes. **B-D** Snapshots of the examples of IARs and DARs showing the regular changes of ATAC-seq and RNA-seq signals in or near CD44 (**B**), DIO2 (**C**) and SULF1 (**D**). **E** The analysis for repeated sequence distributed in IARs and DARs.

**Supplementary Fig. 7.**
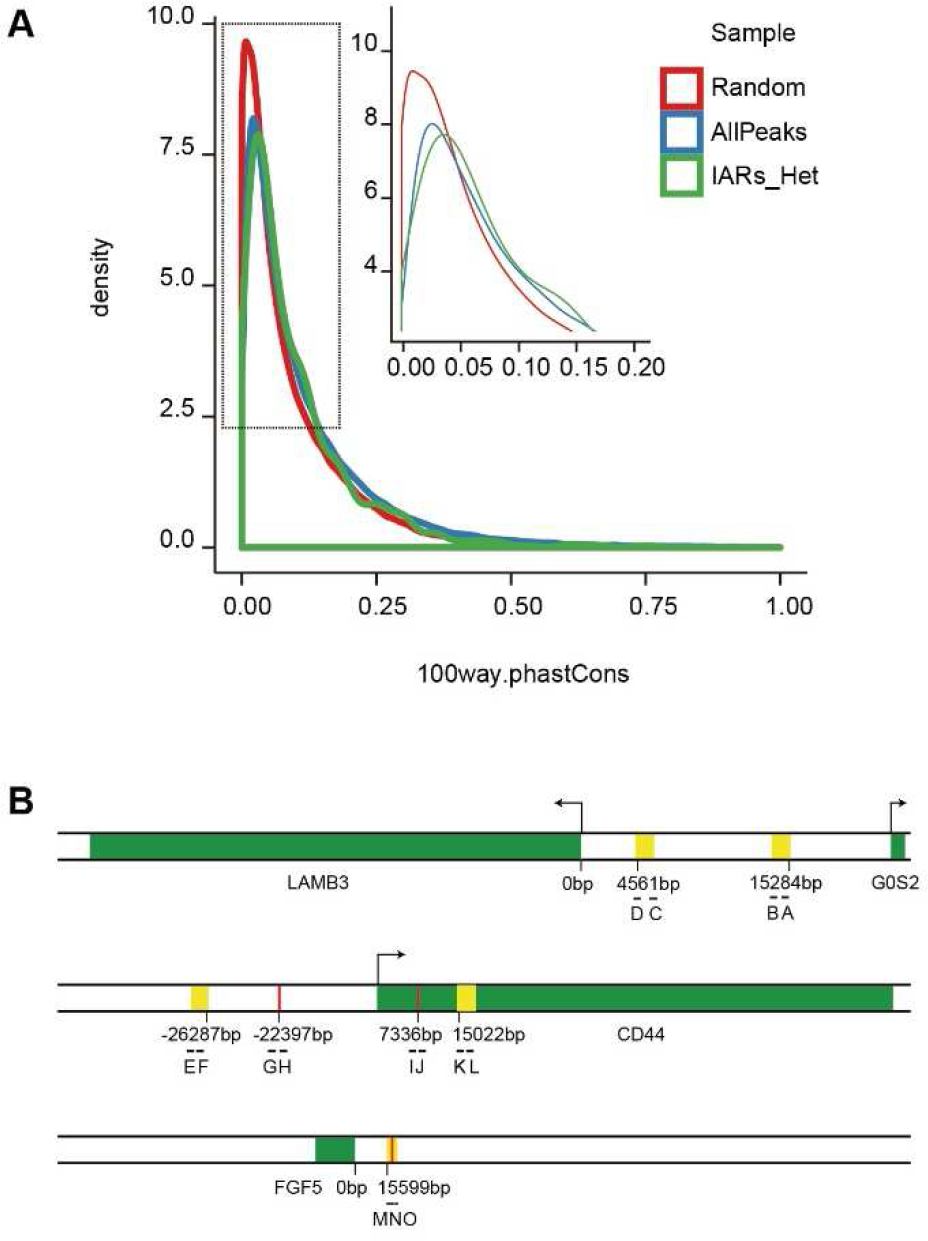
The conservation analysis for IARs and the location information of ChIP-qPCR primers. **A** The analysis for the conservation of IARs in evolution. **B** The relative location of ChIP-qPCR primers used in this research. The red vertical lines represent the predicted AP-1 binding sites, the yellow shadows represent the IARs, the green yellows represent the gene body.

**Supplementary Fig. 8.**
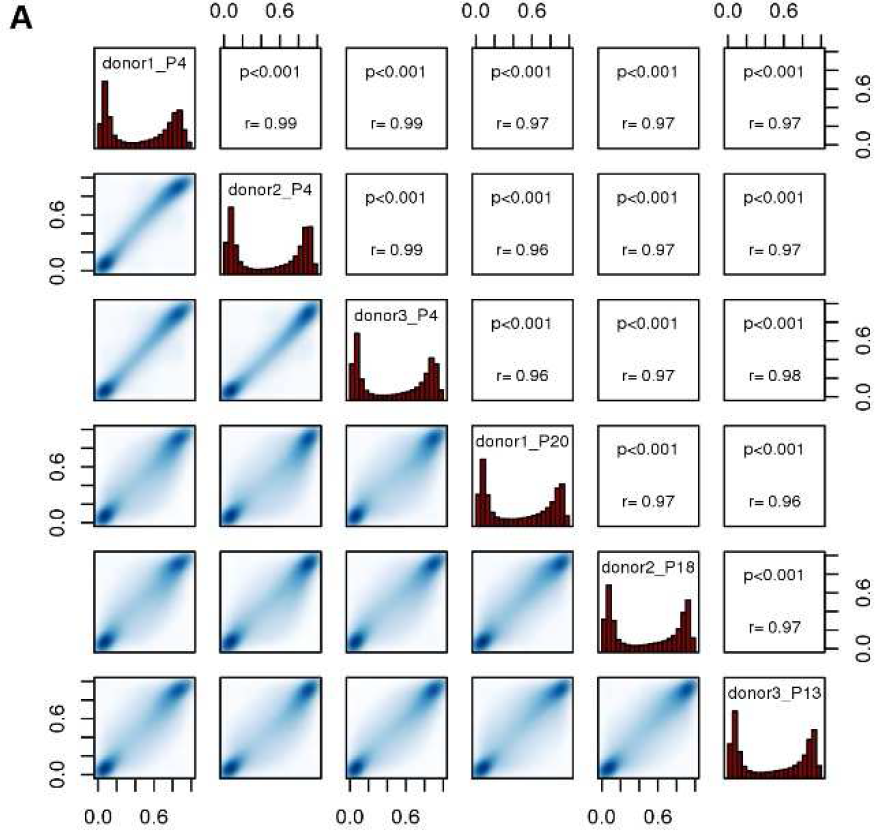
**A** The reliability analysis of DNA methylation showing the strong correlation between different repeated samples.

**Supplementary Fig. 9.**
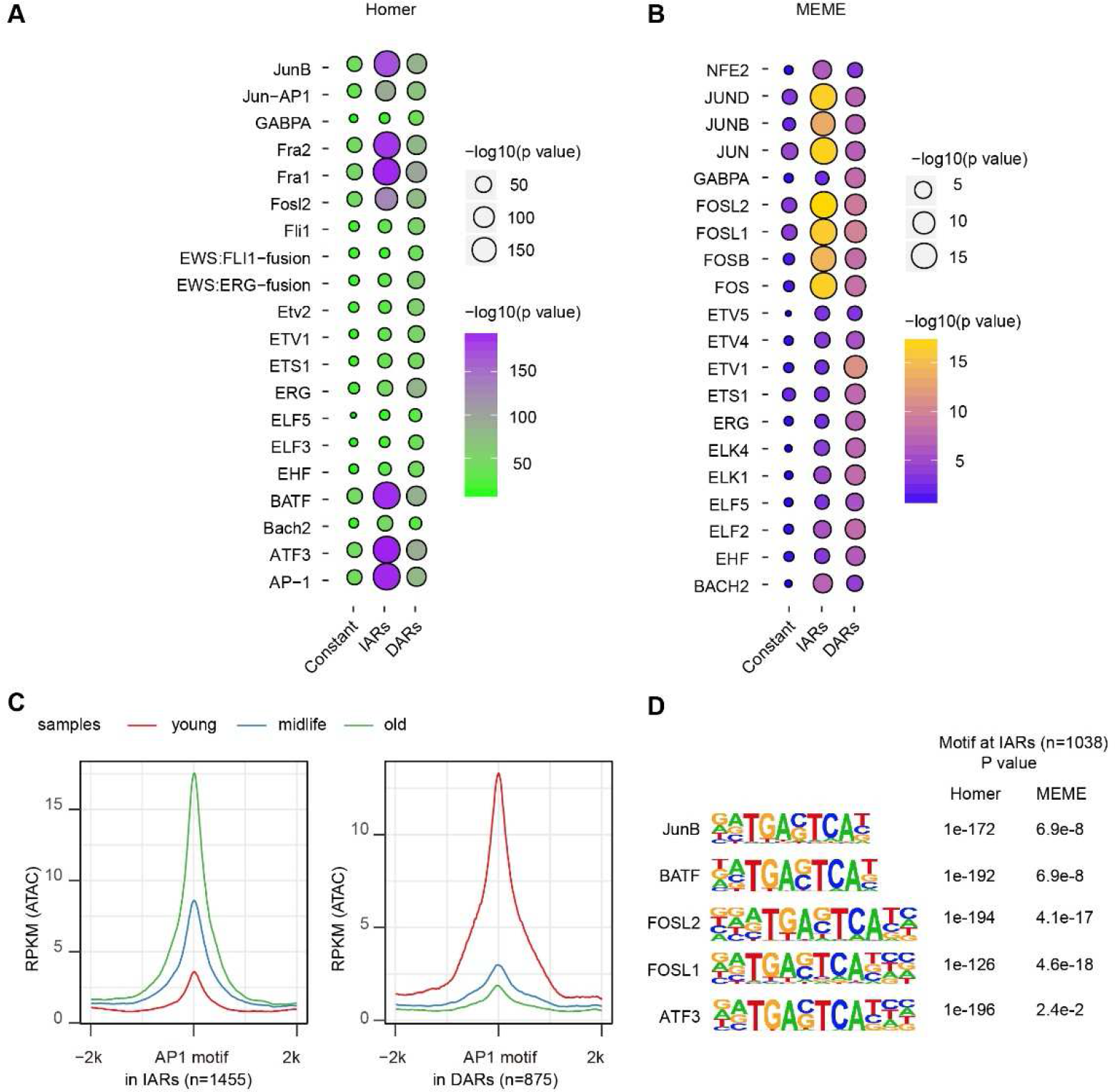
The identification of AP-1 transcription factors in senescence. **A, B** The motifs analysis for transcription factors in IARs and DARs, using Homer (**A**) and MEME (**B**) algorithms respectively. **C** The statistics for RPKM (ATAC) of AP-1 motif in IARs and DARs. **D** The significance of different AP-1 transcription factors in Homer and MEME algorithms.

## Supplementary Table

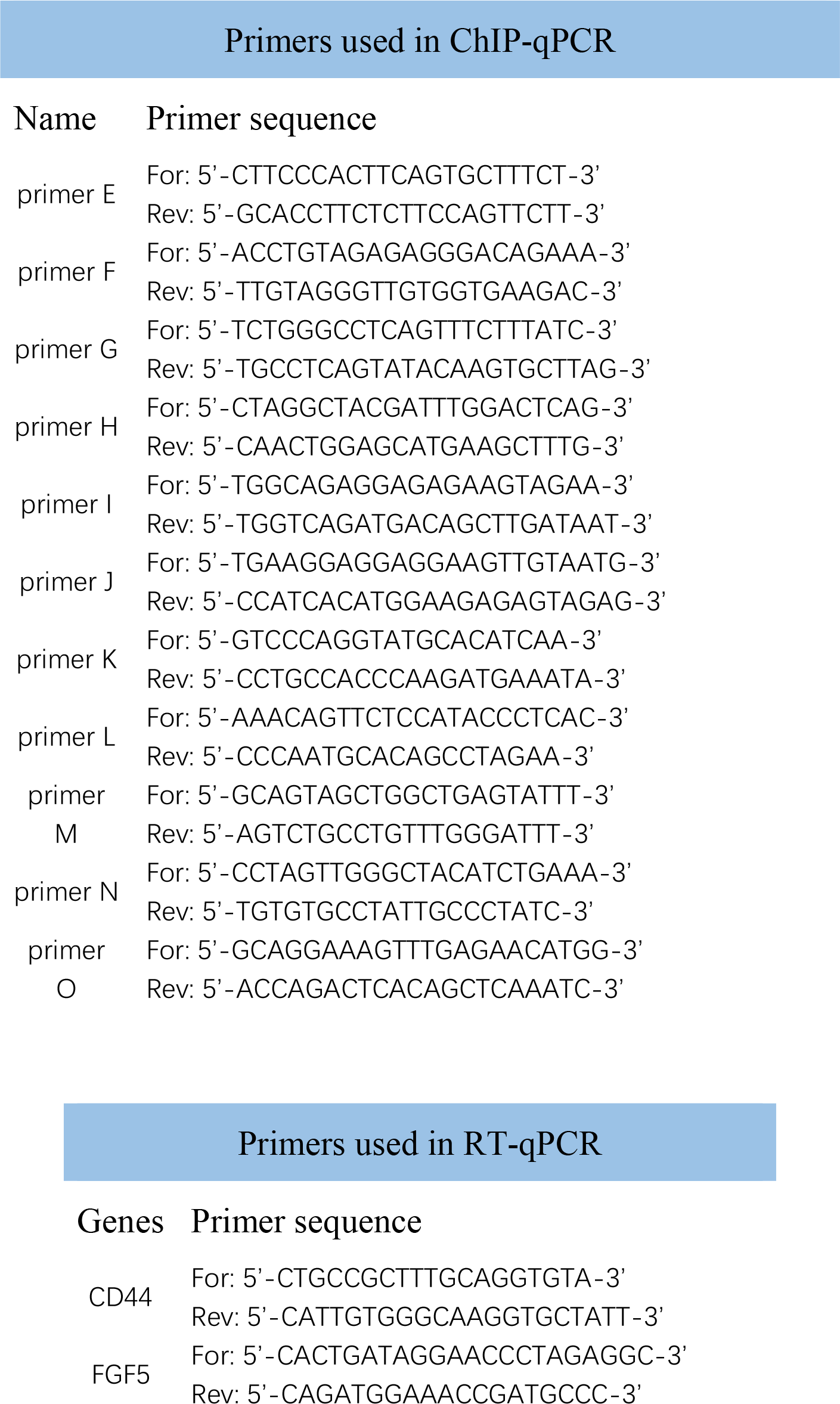

